# Parallel proteomics and phosphoproteomics defines starvation signal specific processes in cell quiescence

**DOI:** 10.1101/2023.08.03.551843

**Authors:** Siyu Sun, Daniel Tranchina, David Gresham

**Affiliations:** Center for Genomics and Systems Biology; Department of Biology, New York University, New York, NY, 10003, USA; Courant Institute of Mathematical Sciences, New York University, New York, NY, 10003, USA

## Abstract

Cells arrest growth and enter a quiescent state upon nutrient deprivation. However, the molecular processes by which cells respond to different starvation signals to regulate exit from the cell division cycle and initiation of quiescence remains poorly understood. To study the role of protein expression and signaling in quiescence we combined temporal profiling of the proteome and phosphoproteome using stable isotope labeling with amino acids in cell culture (SILAC) in *Saccharomyces cerevisiae* (budding yeast). We find that carbon and phosphorus starvation signals activate quiescence through largely distinct remodeling of the proteome and phosphoproteome. However, increased expression of mitochondrial proteins is associated with quiescence establishment in response to both starvation signals. Deletion of the putative quiescence regulator *RIM15*, which encodes a serine-threonine kinase, results in reduced survival of cells starved for phosphorus and nitrogen, but not carbon. However, we identified common protein phosphorylation roles for RIM15 in quiescence that are enriched for RNA metabolism and translation. We also find evidence for RIM15-mediated phosphorylation of some targets, including IGO1, prior to starvation consistent with a functional role for RIM15 in proliferative cells. Finally, our results reveal widespread catabolism of amino acids in response to nitrogen starvation, indicating widespread amino acid recycling via salvage pathways in conditions lacking environmental nitrogen. Our study defines an expanded quiescent proteome and phosphoproteome in yeast, and highlights the multiple coordinated molecular processes at the level of protein expression and phosphorylation that are required for quiescence.

## Introduction

Regulation of cell proliferation and growth in response to extracellular cues such as growth factors, hormones, and nutrients affects development and life span in virtually every biological system (Gray, Petsko, and Johnston 2004). In the absence of stimulatory signals, many cells can enter a reversible quiescent state that is characterized by low metabolic activity and low rates of transcription and protein synthesis. In metazoans, quiescence is induced in response to growth factors and hormones, whereas in unicellular organisms quiescence is primarily initiated in response to nutrient starvation. Quiescence confers distinct physiological properties for cells and is associated with increased stress resistance and recalcitrance to many pharmacologic agents (Sun and Gresham, 2021). The capacity of cells to maintain a viable non-proliferative state for prolonged periods is also referred to as chronological life span (CLS) (Aragon et al. 2008). Despite the universality of the quiescent state, and its potential consequences for aging and disease, the mechanisms regulating entry and maintenance of quiescence remain poorly understood.

Quiescence and CLS have been extensively studied in *Saccharomyces cerevisiae* (budding yeast) primarily in cells grown to stationary phase in nutrient rich medium, in which the starvation signal is undefined. However, as with many microbes, quiescence in yeast can be initiated in response to deprivation of a variety of different nutrients (Lillie and Pringle 1980; Gresham et al. 2011; Klosinska et al. 2011; Yanagida 2009; Sun et al. 2020). Starvation for essential nutrients other than carbon, including nitrogen, phosphorus, and sulfur result in many of the same characteristic features including cell cycle arrest as unbudded cells, thickened cell walls, increased stress resistance, and an accumulation of storage carbohydrates (Lillie and Pringle 1980; Klosinska et al. 2011; Schulze et al. 1996). The ability to effectively initiate, maintain, and exit quiescence confers a significant selective advantage across diverse environments resulting in evolutionary selection for effective cellular quiescence (O’Farrell 2011). However, how these distinct nutrient starvation signals converge on the same cellular response is not well understood.

Cell cycle exit and establishment of cell quiescence is controlled by a signaling cascade that is initiated in response to external signals. In budding yeast, RIM15, an evolutionary conserved serine/threonine kinase has been identified as a downstream target regulated by multiple nutrient responsive pathways including Target Of Rapamycin Complex 1 (TORC1), protein kinase A (PKA), or PHO80-PHO85 (PHO) (Swinnen et al. 2006; Sarkar et al. 2014). The molecular pathways linking RIM15 to distal readouts including the transcription of specific nutrient-regulated genes, trehalose and glycogen accumulation, extension of CLS, and induction of autophagy have only been partially characterized in stationary phase cultures grown in nutrient rich conditions (Pedruzzi 2000; Wei et al. 2008; Yorimitsu et al. 2007).

Increased protein turnover during the transition from proliferation to quiescence is essential for protein homeostasis and ensuring viability and long term survival (Marguerat et al. 2012; Zakrajšek, Raspor, and Jamnik 2011). Whether cells remodel their proteome differently according to different nutrient deprivation cues and use the same mechanisms for mediating protein homeostasis during this transition has not been determined. Recently, we found that RIM15 genetically interacts with proteostasis genes that control protein translation and degradation (Sun et al. 2020). We hypothesized that RIM15 integrates information about distinct nutrient signals to coordinate protein expression in quiescent cells. To test this hypothesis, we used stable isotope labeling by amino acid in cell culture (SILAC) and mass spectrometry in prototrophic yeast strains to define the proteome and phosphoproteome during quiescence in response to defined starvation signals: carbon, nitrogen, and phosphorus. In wildtype cells we observed large-scale proteome remodeling in response to different starvation signals and identified both a common starvation response signature as well as starvation signal specific changes in global protein expression. Increased mitochondrial protein expression is a common response regardless of nutrient starvation signal. We found that the phosphoproteome undergoes an even more dramatic remodeling compared to proteome. Deletion of (*RIM15Δ0*) differentially impacts proteome and phosphoproteome dynamics during quiescence entry under different starvations. Consistent with genetic interaction mapping we identified potential new phosphorylation targets of RIM15 that are associated with protein synthesis and degradation. Overall, our results are consistent with a combination of common and nutrient-specific processes mediating quiescence and redundant pathways that operate in parallel to RIM15 to regulate quiescence.

## Results

### Prototrophic yeast strains efficiently incorporate exogenous lysine and arginine

SILAC is a method for labeling proteins *in vivo* using isotope-containing amino acids provided in the culturing media. Engineered auxotrophic yeast strains that cannot synthesize lysine and arginine efficiently import and assimilate exogenously provided amino acids containing different isotopes of carbon or nitrogen. However, auxotrophic cells exhibit impaired starvation signal processing, and concomitant low viability in response to starvation, precluding their use in the study of quiescence (Boer, Amini, and Botstein 2008; Gresham et al. 2011). It has previously been demonstrated that in prototrophic yeast strains proteins can be efficiently labeled with isotopic lysine (lys-8), likely because amino acid incorporation from the culture medium is less energetically demanding than *de novo* biosynthesis (Fröhlich, Christiano, and Walther 2013). To test whether SILAC can be performed with prototrophic strains of different genotypes, we first estimated labeling efficiency by growing different yeast strains in the same SILAC medium (**Figure 1A**). We found that both prototrophic and auxotrophic wildtype (WT) strains and a prototrophic *RIM15* deletion mutant (*RIM15Δ0*) have similar doubling times in standard SILAC media (i.e. synthetic complete (SC) media containing 111mM glucose) with supplemented heavy (^13^C_6_^15^N_2_-lysine [Lys-8], ^13^C_6_^15^N_4_-arginine [Arg-10]) or medium (4,4,5,5-D4 lysine [Lys-4], ^13^C_6_-arginine [Arg-6]) isotopic labels. We extracted proteins from all three strains (WT prototroph, *lys2-arg4-* auxotrophs, and *RIM15Δ0* prototroph) and performed LC-MS/MS. For all strains the label incorporation for each protein (isotope intensity/total intensity) was over 98% (**Figure 1B**). No significant reduction in incorporation level was observed for either WT or *RIM15Δ0* prototrophic cells in comparison to auxotrophic cells. Moreover, the proteome of prototrophic and auxotrophic wildtype cells are highly correlated in both medium (Lys-4 and Arg-6) SILAC media (*r* = 0.93) and heavy (Lys-8 and Arg-10) SILAC media (*r* = 0.95) (**Figure S1A**). By contrast, the correlation between *WT* and *RIM15Δ0* proteomes is considerably reduced (*r* = 0.8) (**Figure S1B** and **Figure S1C**), indicating that genotype is the primary source of difference in the proteomes. Together, our results confirm the suitability of SILAC to study the proteome of prototrophic cells.

**Figure 1.**
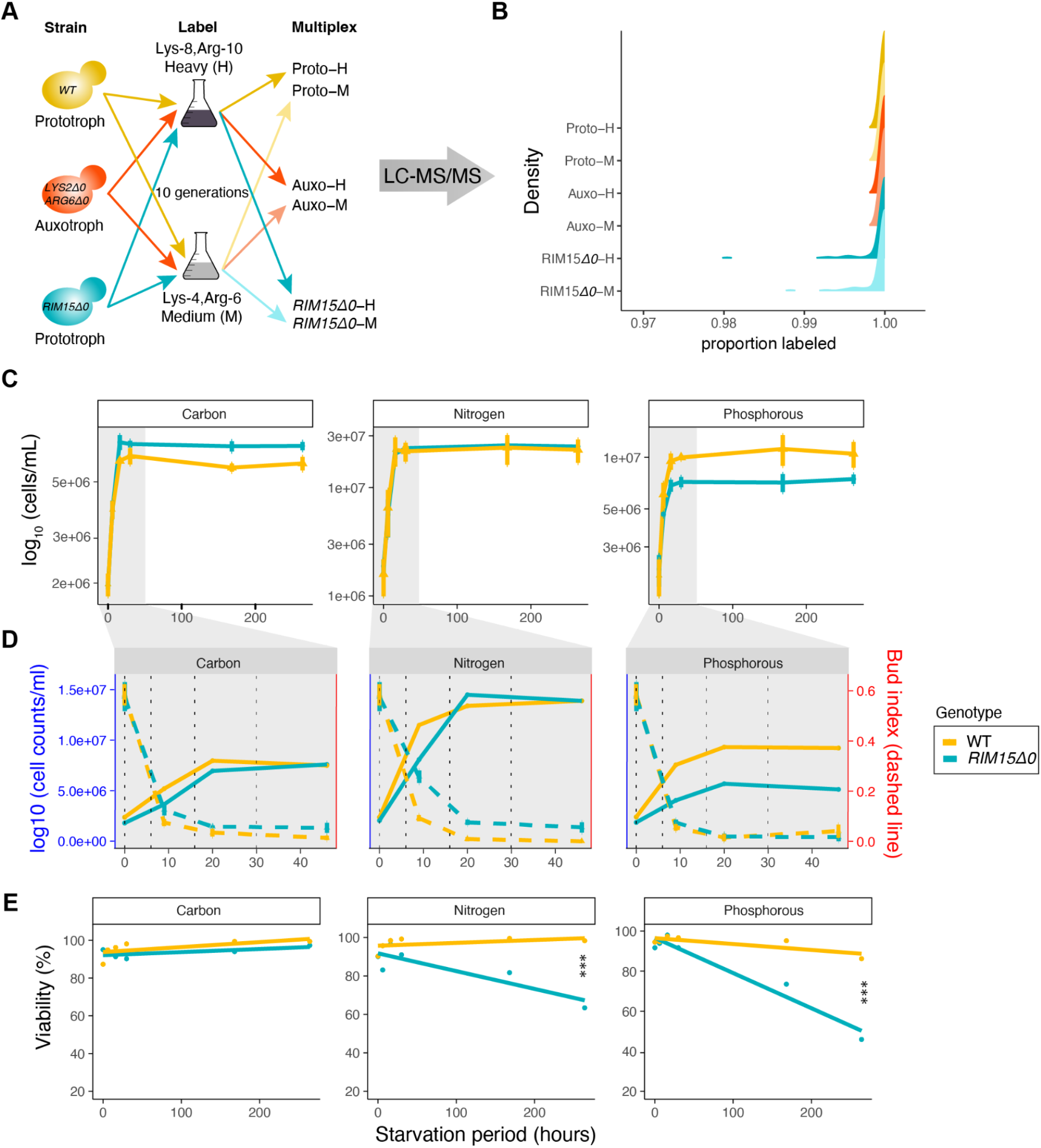
The effect of *RIM15* deletion on cell cycle arrest and survival in different nutrient starvation conditions. **A)** Experimental design for testing amino acid incorporation in prototrophic and auxotrophic strains. Prototrophic wild type (Proto-), auxotrophic (*LYS2Δ0 ARG6Δ0*) wildtype (Auxo-), and prototrophic mutant (*RIM15Δ0*) cells were grown in either Heavy (^13^C_6_^15^N_2_-lysine [Lys-8], ^13^C_6_^15^N_4_-arginine [Arg-10]) or Medium (D4-lysine [Lys-4], ^13^C_6_-arginine [Arg-6]) isotope-supplemented SILAC media. Cells were collected for LC-MS/MS analysis following 10 generations of growth. **B)** Incorporation levels of heavy (Lys-8, Arg-10) and medium (Lys-4, Arg-6) isotope-containing proteins for the auxotrophic and prototrophic *S. cerevisiae* S288c strain. **C)** Growth dynamics of prototrophic wild type (yellow solid line) and *RIM15Δ0* (blue solid line) cells following transfer to three nutrient starvation media: carbon - 0.06% (3.35mM) glucose, Nitrogen - 0mM nitrogen, Phosphorous - 0mM phosphorus (n = 3) with addition of 100mM lysine and 100mM arginine. T = 0 is the time at which cells were transferred into the corresponding nutrient-depleted condition from mid-log growth in rich media. **D)** Growth (solid line) and bud-index (dashed line) for both wildtype (yellow) and *RIM15Δ0* (blue) cells during the first 46 hours post nutrient removal. *RIM15Δ0* cells cease proliferation and arrest as unbudded cells at the same time as wild type cells. **E)** Viability of wildtype and *RIM15Δ0 cells* measured using PI/Syto9 staining and flow cytometry. Statistical significance of slope difference between survival curves of wildtype and *RIM15Δ0* is shown. *** indicates Bonferroni corrected p-value < 0.001.

### Loss of RIM15 reduces survival of nitrogen and phosphorus starvation, but not carbon starvation

To test the cell biological response of prototrophic WT and *RIM15Δ0* cells in different starvation conditions we monitored growth, bud index, and survival when cells were transferred from rich medium to different defined starvation media. In preliminary experiments we found that acute starvation for carbon resulted in a sudden cessation of growth whereas cells starved for phosphorus or nitrogen continued to grow for at least two population doublings (i.e. an average of two additional mitotic cell divisions) likely as a result of internal phosphorus and nitrogen stores. Therefore, to equalize population growth and arrest dynamics we provided a small amount of carbon in carbon starvation media (3.35mM glucose) whereas the nutrient was completely absent in nitrogen and phosphorus starvation media. Upon nutrient depletion, both WT and *RIM15Δ0* cells doubled 2-4 times (depending on the starvation condition) before arresting growth (**Figure 1C**). Bud index analysis showed that in the absence of RIM15, cells still exhibit uniform cell cycle arrest in G1 (**Figure 1D**). As RIM15 has been proposed to be a key regulator for quiescence establishment in undefined starvation conditions (Bontron et al. 2013), we tested for survival defects. In the absence of RIM15 long-term survival in response to nitrogen and phosphorus starvation was significantly reduced; however, no significant defect was observed in response to carbon starvation (**Figure 1E**). These results indicate that RIM15 has distinct roles in conferring long term survival in response to different starvation signals, but does not play a significant role in starvation induced cell cycle exit.

### Multiplexed quantitative analysis of the proteome and phosphoproteome during quiescence initiation

We next sought to leverage SILAC labeling to identify protein expression and phosphorylation states during quiescence establishment in response to different nutritional starvations. Triplicate cell populations were cultured with isotopically labeled lysine and arginine to enable multiplexing cells with different genotypes and to eliminate batch effects when quantifying proteomes from different genotypes across time. Specifically, quiescent samples were labeled with either heavy or medium amino acids and a universal sample of unlabeled (light) protein from log phase WT cells was generated and mixed with all experimental samples for quality control and normalization of each mass spectrometry run (**Figure S1D**). Samples were acquired in log-phase (0 hours), and then at 6, 16, and 30 hours post nutrient depletion. Samples were lysed, pooled, digested, and analyzed by LC-MS/MS (**Methods**). A total of 5% of the pooled sample was used for whole proteome analysis, and the remaining 95% was subjected to further fractionation using titanium oxide columns for phosphoproteome profiling. In total, we quantified 1,277 proteins (**Table S1)** and 1,472 phosphorylation events (**Table S2)** at a false discovery rate (FDR) <1%, corresponding to approximately 25% of the total yeast proteome.

We first assessed technical variation by determining the ratio for each labeled protein relative to the unlabeled universal sample. Surprisingly, we observed a significant discrepancy between the distributions of heavy/light and medium/light ratios specifically in nitrogen starvation conditions. In all biological replicates, we observed a marked reduction in detected proteins labeled with heavy isotopes (**Figure S1E**). To determine the source of this unexpected reduction in protein from nitrogen starved cells, and exclude the possibility of technical issues associated with mass spectrometry analysis, we performed LC-MS/MS analysis of samples from nitrogen starvation conditions that were cultured in heavy labeling conditions (Lys-8 and Arg-10) without subsequent mixing with samples containing light or medium label. We first checked whether samples from later time points had fewer MS scans, which is a known factor that results in limited protein detection. Interestingly, similar numbers of MS scans were found for both MS1 and MS2 (MS/MS) across three tested time points (**Figure S1F**). However, fewer MS2 scans were identified as a function of time, leading to an increase in peptides that cannot be matched with the reference database (**Figure S1F and Figure S1G**). Inspection of the raw spectra for several peptides at different time points following nitrogen starvation identified additional peaks that were not detected prior to starvation (**Figure S1H**). These observations indicate that the reduced number of isotopically labeled proteins identified in nitrogen starvation conditions is a result of incorporation of isotopic labels into amino acids other than lysine and arginine. Thus, in the absence of nitrogen, cells appear to catabolize the heavy SILAC amino acids (Lys-8 and Arg-10) to enable synthesis of other amino acids. Therefore, SILAC is not an appropriate experimental method for quantitatively assaying proteome remodeling during quiescence in response to nitrogen starvation and these samples were excluded from subsequent analyses.

We then sought to understand sources of variation in proteome and phosphoproteome data for cells starved for either carbon and phosphorous. To differentiate technical variation from biological variation, we compared replicate data at T=0 hour and T=30 hour for both whole proteome and phosphoproteome. We found that technical variation is randomly distributed for both proteome (**Figure S2A** left panel) and phosphoproteome (**Figure S2B** left panel) data with phosphoproteome data being more variable than the whole proteome data, likely due to the transient nature of phosphorylation. By contrast, comparison between proliferating and quiescent cells reveal significant biological variation that is reproducible between replicates for both proteome (**Figure S2A** right panel) and phosphoproteome **(Figure S2B** right panel) data. Principal-component analysis (PCA) of protein (**Figure S2C**) and phosphopeptide (**Figure S2D**) abundances further support biological variation as the main source of variance in the data.

### Mitochondrial protein expression is increased in response to carbon and phosphorus starvation

We quantified the dynamics of protein expression in wildtype cells starved for either carbon or phosphorus. Expression dynamics of several proteins measured by mass spectrometry are in agreement with results from a previous study using GFP fusion proteins (Davidson et al. 2011). In total, 44 of 125 mitochondrial proteins previously reported to increase in expression in rich media starvation conditions (Davidson et al. 2011; Allen et al. 2006) were also identified in the proteome of carbon-starved cells. We found that 37 of the 44 proteins systematically increase in expression as a function of time (**Figure 2A**). Notably, 25 out of 38 quiescence specific upregulated proteins (Davidson et al. 2011), which includes 9 proteins that are not localized to the mitochondria, are also found in carbon starved quiescent cells in our study (**Figure 2B**). Overall, our results using mass spectrometry are consistent with previous findings using a distinct method of proteome analysis, thereby validating our approach.

**Figure 2.**
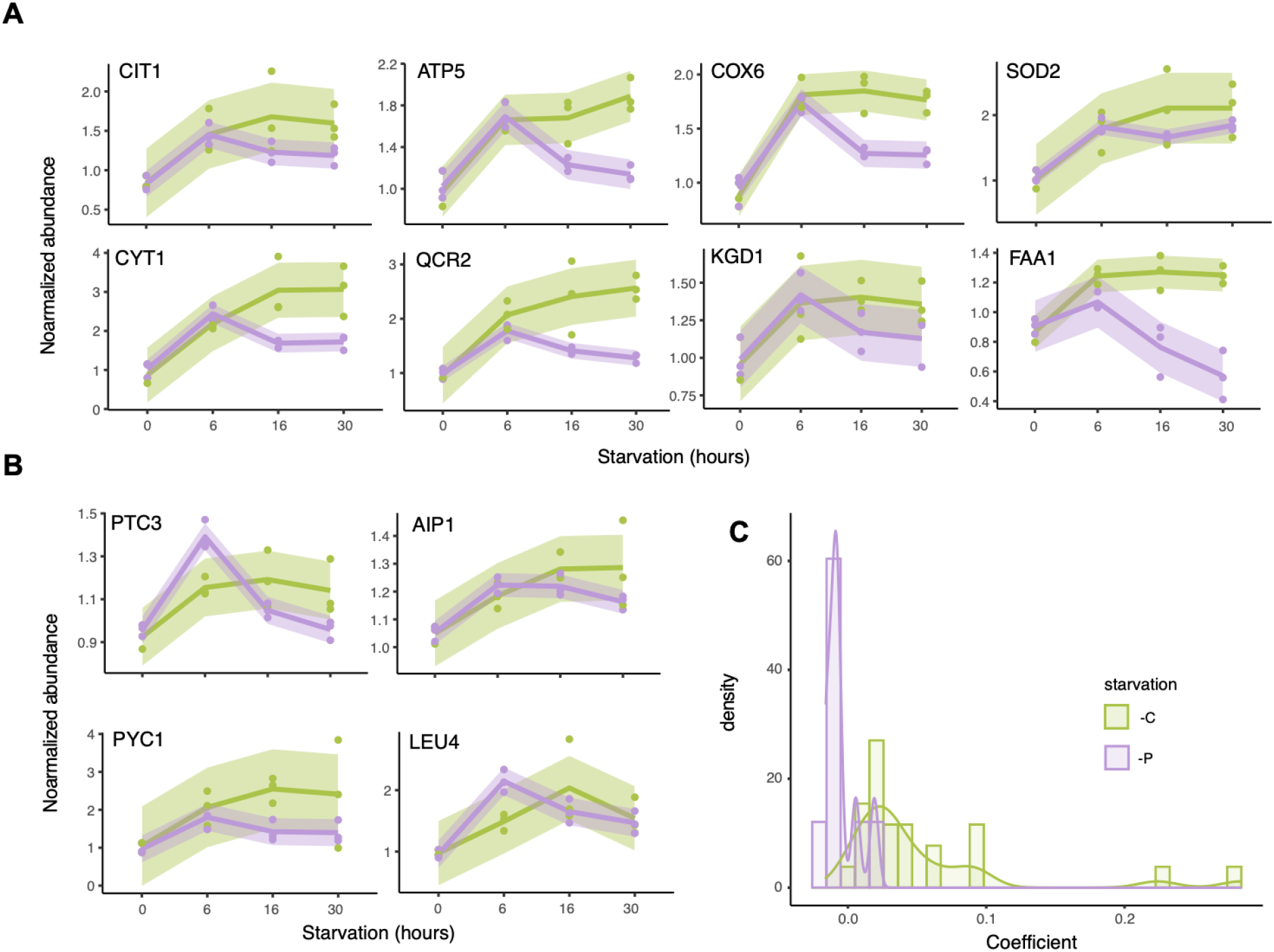
Mitochondrial proteins increase in expression in response to both carbon and phosphorus starvation. **A)** Examples of known quiescence-specific upregulated mitochondrial proteins that are localized to mitochondria (Davidson et al. 2011) in carbon (green line) and phosphorus (purple line) starvation induced quiescent wild type cells. Solid lines connect the mean expression at each timepoint. Shading indicates standard error. **B)** Representative examples of non-mitochondrial proteins that are upregulated in quiescence (Davidson et al. 2011) in carbon (green line) and phosphorus (purple line) starvation induced quiescent wild type cells. **C)** Distribution of coefficients from linear regression models fit to proteins quantified by mass spectrometry in wild type cells starved for carbon or phosphorus that have previously identified as being upregulated in quiescence in rich media starvation conditions using a GFP library (Davidson et al. 2011).

We find that the majority of these previously identified quiescence-related proteins also increase in expression in response to phosphorus starvation. However, in contrast to their monotonic increase in carbon starvation conditions, the expression of these proteins peaks 6 hours after phosphorus starvation and is subsequently attenuated (**Figure 2A** and **Figure 2B**). To quantify the increase in expression following initiation of starvation we fit linear models to the expression of each protein. The transient increase in expression in phosphorus results in a reduced distribution of linear model coefficients (i.e. slope) compared with carbon starvation **(Figure 2C)**. The upregulation of mitochondrial proteins in response to both defined glucose and phosphorus starvation, as well as rich media starvation (Davidson et al. 2011; Allen et al. 2006; Aragon et al. 2008), points to an important role for increased mitochondrial function in response to at least two different quiescence-inducing signals.

### Identifying protein correlation networks in quiescence

We sought to systematically model the dynamics of all protein expression changes during the initiation of quiescence in response to distinct starvation signals. We adopted a two-step computational pipeline (**Figure 3A**) (Tan et al. 2017): 1) we identified proteins that significantly changed in abundance between any two time points, genotype or starvation condition using three-way ANOVA, which resulted in identification of 1,067 differentially expressed (DE) proteins (FDR < 0.05) among the 1,277 measured proteins (**Table S1**), 2) we applied Weighted Gene Correlation Network Analysis (WGCNA) to DE proteins to identify modules defined by correlated protein expression dynamics. Modules were then tested for association with study variables (i.e. starvation condition, genotype, and time) to identify protein functions that vary in behavior as a function of experimental variables. Using WGCNA we grouped the majority of the DE proteins into six whole proteome modules (WPMs) that exhibit distinct expression patterns across genotypes, nutrient starvation, and time (**Figure 3B**).

**Figure 3.**
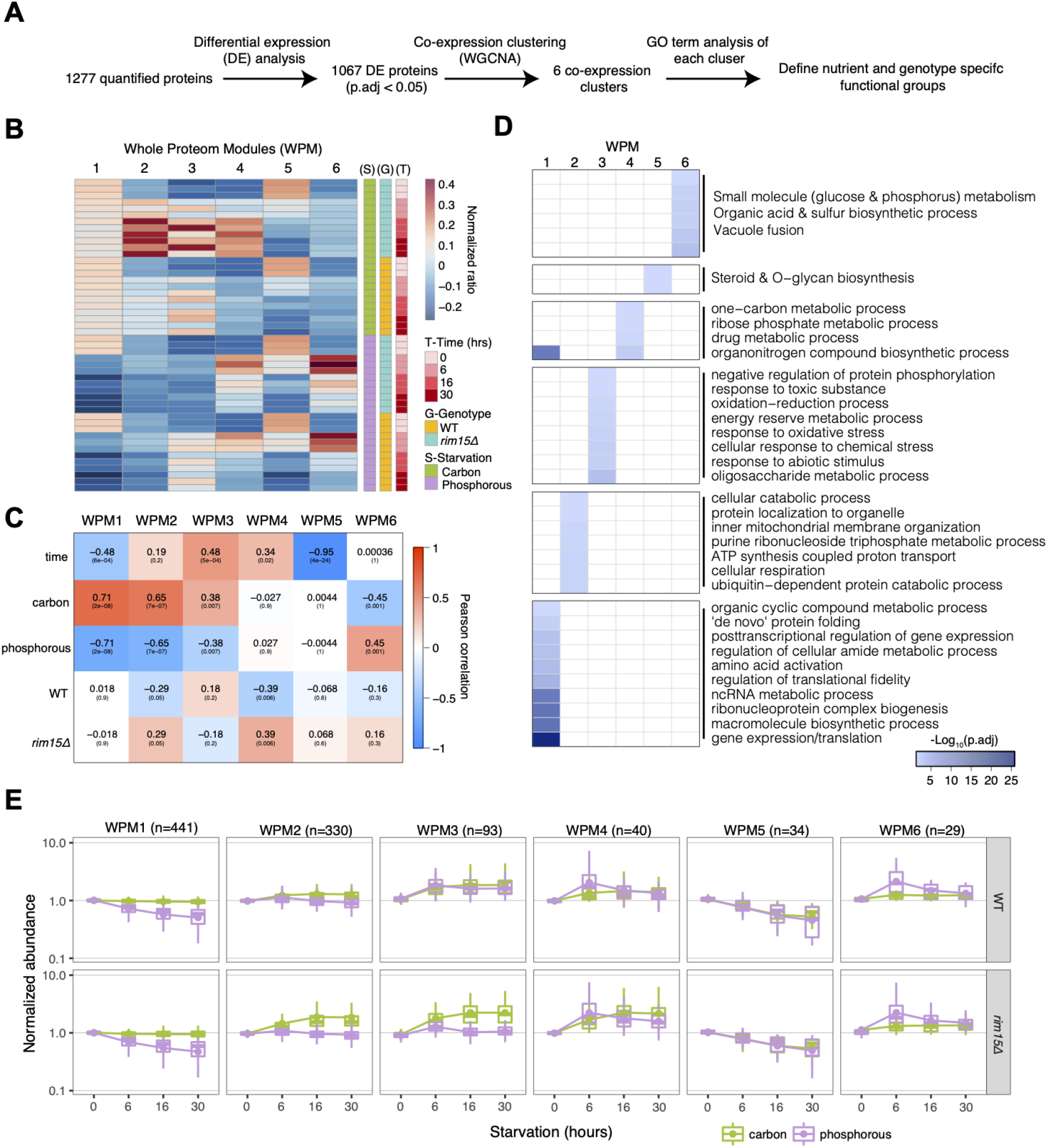
Temporal expression profiling of the yeast proteome reveals correlated expression of whole proteome modules (WPMs) during initiation of quiescence. **A)** Overview of computational workflow. **B)** Expression of eigengenes for each whole proteome module (WPM). Each column corresponds to a whole proteome module and each row corresponds to an individual sample. Each experimental condition was measured in biological triplicate. **C)** WPM to experimental variable relationships as determined by Pearson correlation coefficient. Statistical significance for each coefficient is indicated. **D)** Functional annotation of WPMs using Gene Ontology and KEGG pathway annotation enrichment (FDR < 0.05). **E)** Expression distribution for all proteins in each WPM across time for wildtype and *RIM15Δ0* cells in phosphorus and carbon starvation.

We used correlation between WPMs and experimental variables to quantitatively define the strength of their relationships (**Figure 3C**) and annotated WPMs using Gene Ontologies and KEGG pathways (**Figure 3D**). Employing this approach we find that the specific nutrient starvation signal is the primary source of variation in protein expression. WPM1, the largest module (441 proteins), is significantly and gradually repressed in phosphorus starvation conditions but exhibits minimal variance in carbon starvation **(Figure 3C** and **Figure 3E**). Functional annotation of proteins in WPM1 reveals enrichment for different aspects of protein homeostasis regulation including post-transcriptional modification, tRNA charging, translation fidelity, and protein folding (**Figure 3D**). Mitochondrial functions are enriched for proteins found in WPM2 (330 proteins), including those discussed above (**Figure 2**), which peaks in expression at 6 hours in both starvation conditions. Expression of WPM2 subsequently remains stable throughout the time course in carbon starvation but decreases at later time points in phosphorus starvation (**Figure 3E**) as observed for individual proteins (**Figure 2A**).

Proteins in WPM3 (93 proteins), enriched for stress-related functions (**Figure 3D**), and WPM6 (29 proteins), enriched for small molecular metabolism and vacuole fusion (**Figure 3D**), increase in expression in both starvation conditions but exhibit different dynamics. Proteins in WPM6 are upregulated after 6 hours of starvation and subsequently decline in expression in phosphorus starvation whereas these proteins are gradually upregulated throughout the carbon starvation and ultimately reach a similar expression to the phosphorus starvation condition (**Figure 3E**). Proteins in WPM3 rapidly increase in expression in both starvation conditions, but tend to continue to increase modestly in carbon starvation but not phosphorus starvation.

By contrast, WPM5 (34 proteins) exhibits strikingly similar behavior in carbon and phosphorus starvation (**Figure 3E**), with significant repression over time (**Figure 3C**). WPM5 is enriched for proteins involved in steroid and O-glycan biosynthesis pathways (**Figure 3D**). The consistent down regulation of proteins in WPM5 regardless of starvation signal suggests a common strategy for energy conservation by minimizing specific biosynthetic activities to do with cell membrane and cell wall growth.

The most significant effect of the loss of RIM15 is observed for WPM2, WPM3, and WPM4 (**Figure 3C** and **Figure S3A**) as it appears to result in an amplified protein expression response relative to wildtype specifically in glucose starvation conditions (**Figure 3E** and **Figure S3A**). Thus the primary effect of RIM15 on the proteome of carbon starved cells appears to be in regulating the magnitude of protein expression changes perhaps by dampening overall protein production. By contrast, in response to phosphorus starvation, *RIM15Δ0* cells fail to increase expression of proteins in WPM3 (**Figure S3A**). As WPM3 is enriched for proteins involved in multiple stress responses (**Figure 3D**), this failed induction specially in response to phosphorus starvation may contribute to the reduced survival of the *RIM15Δ0* in phosphorus starvation conditions (**Figure 1E**).

Overall, the majority of proteome remodeling occurs during the initial starvation period and is largely stable from 16 to 30 hours after starvation (**Figure 3E**). These expression dynamics are consistent with the behavior of the population (**Figure 1C**); in the first two time points (T = 0 hours and T = 6 hours) the cells sense the starvation signals, complete the last round of cell division, and prepare to enter quiescence. By the second two timepoints (T = 16 hours and T = 30 hours) the cells have arrested growth and appear to fine tune protein expression to maintain minimal cellular activities.

### RIM15 coordinates biosynthetic pathways and mitochondrial metabolism in quiescence

To quantify individual protein expression dynamics during quiescence initiation that are regulated by RIM15 we applied analysis of covariance (ANCOVA) (**Methods**). We detected 298 proteins in carbon starvation and 82 proteins in phosphorus starvation that have different dynamics as a function of *RIM15* genotype (p.adj < 0.05) (**Figure S3B**). Consistent with the results from WGCNA analysis, the majority (75%) of differentially expressed proteins are increased in expression in the absence of RIM15 under carbon starvation, suggesting a primarily repressive function for RIM15.

In contrast to carbon starvation, a similar number of proteins are found to be up-(n = 43) and down-(n = 39) regulated in phosphorus starvation in the absence of RIM15. Although far fewer proteins (i.e. 39 vs 298) are found to be repressed by RIM15 under phosphorus starvation, a significant number of them (18) are common to both carbon and phosphorus starvation (p.adj < 0.01, hypergeometric test) (**Figure S3D**). The majority of these proteins are enzymes involved in carboxylic acid catabolism (ASC2, LYS21), mitochondrial metabolism (IDP1, ADH3, MMF1) and amino acid synthesis (a total of 8 proteins). Proteins that function in different aspects of protein homeostasis also show evidence of being regulated by RIM15, such as protein folding chaperons (HSP60), ER-associated proteins (CPR5 and PDI1), and protein transport (SEC17). This points to a common repressive role for expression of specific proteins by RIM15 in response to carbon and phosphorous starvation.

We ranked proteins by fold change and applied Gene Set Enrichment Analysis (GSEA) (**Methods and Materials**). Membrane transport proteins that are essential for energy production and synthesis in mitochondria have increased expression in the absence of RIM15 under carbon starvation (**Figure S3C**), again suggesting that RIM15 functions to dampen their expression. By contrast, ribosomal functions are enriched for proteins that have increased expression in WT relative to *RIM15Δ0* cells in carbon starvation indicative of a failure to maintain translation activity in the absence of RIM15. Phosphorus starved *RIM15Δ0* cells fail to upregulate mitochondrial proteins (**Figure S3C)** likely contributing to the poor viability of these cells (**Figure 1C**).

To assess whether upregulated proteins are essential for quiescence, we compared protein expression dynamics with survival data for the corresponding gene deletion upon carbon or phosphorus starvation (Sun et al. 2020). Overall, there is no relationship between protein expression and survival of the gene deletion mutant in quiescence (**Figure S3E**). However, there is clear evidence for the genetic importance of the many proteins that increase in expression in quiescence. A total of 11 of 51 proteins (∼22%) that are upregulated in carbon starvation (p.adj < 0.05) are genetically required for survival (p.adj < 0.05) (**Table S3**) and 13 of 39 proteins (∼33%) that are upregulated in phosphorus starvation are required for survival (**Table S4**).

### RIM15 regulates similar phosphorylation events in response to different starvation signals

To identify protein phosphorylation targets of RIM15 we performed differential protein phosphorylation analysis at each time point (**Figure S4A**). The known RIM15 phosphorylation target, IGO1_64 (Talarek et al. 2010) has reduced phosphorylation in proliferating cells lacking RIM15 (**Figure 4A**), and this trend is maintained throughout the quiescent period regardless of the starvation signal (**Figure 4B**). This observation is consistent with RIM15 playing an active role in proliferating cells in addition to its function in regulating quiescence (Sarkar et al. 2014). Phosphorylation of TPS2, the phosphatase subunit of the trehalose-6-phosphate synthase, is significantly reduced in a the absence of RIM15 suggesting it is a direct target consistent with a role for RIM15 in regulating storage carbohydrate metabolism. Motif analysis identified the same major RIM15 protein phosphorylation site (Serine/Threonine) in both carbon or phosphorus starvation, with minor differences in the surrounding context (**Figure 4C**).

**Figure 4.**
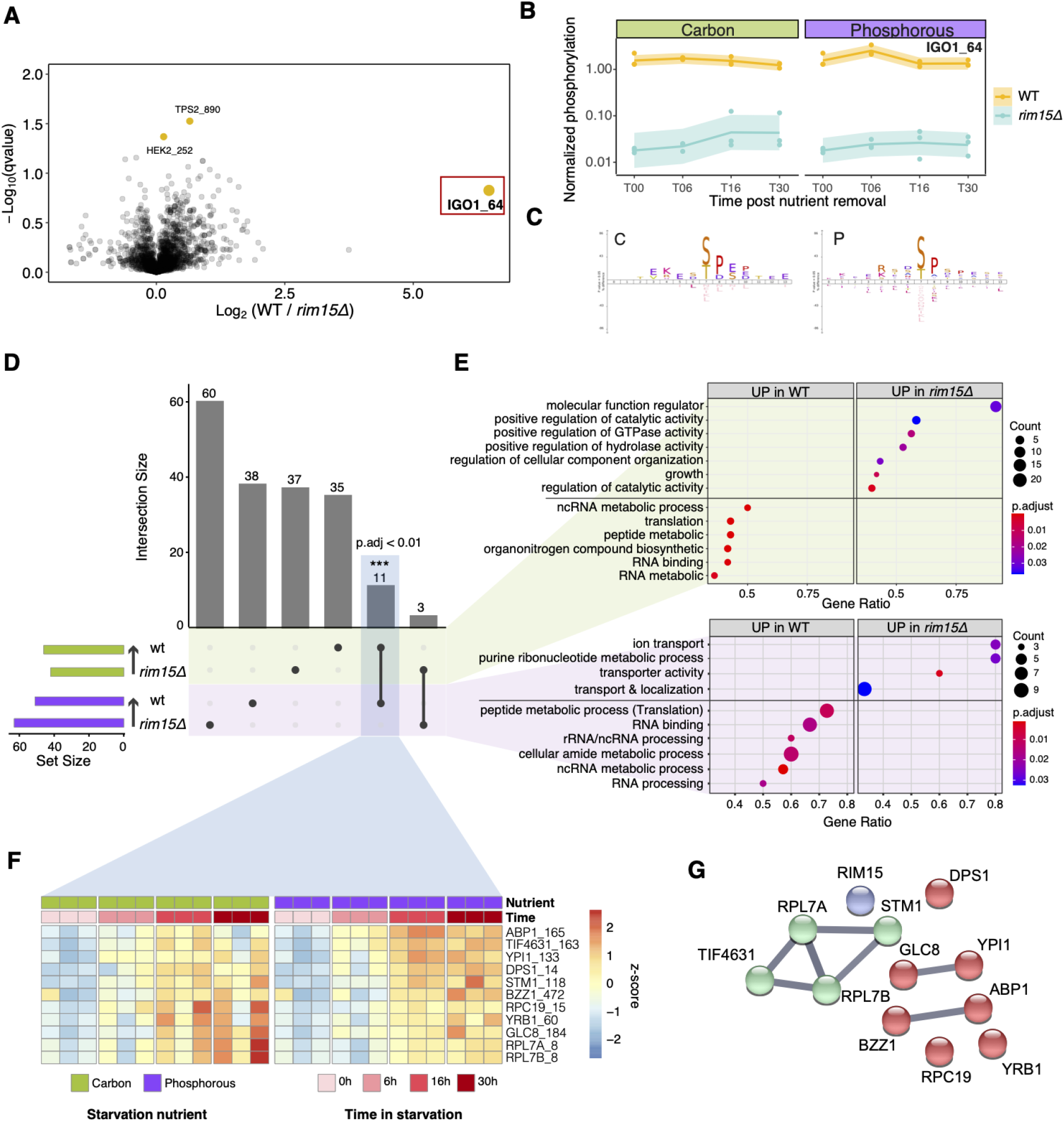
RIM15 protein phosphorylation targets in proliferating and quiescent cells. **A)** Differential protein phosphorylation events in proliferating WT and *RIM15Δ0* cells (**methods and materials**). The previously reported IGO1_64 site is highlighted. **B)** Dynamics of IGO1_64 phosphorylation site in WT and *RIM15Δ0* cells in carbon and phosphorous starvation. **C)** Phosphorylation motif of RIM15 targets in carbon and phosphorous starvation conditions. **D)** Common and unique protein phosphorylation events in WT and *RIM15Δ0* cells. **E)** GSEA of differential phosphorylation events between WT and *RIM15Δ0* cells in response to carbon (green) and phosphorus starvation (purple). **F)** Increased protein phosphorylation of 11 common targets in wildtype cells in both carbon and phosphorus starvation log_2_(WT/*RIM15Δ0*) for each timepoint and replica (each column) during quiescence. **G)** Known functional relationships among 11 proteins whose phosphorylation state in quiescence is dependent on RIM15.

To identify phosphorylation targets of RIM15 in quiescence we used analysis of covariance (ANCOVA) (**Methods**). A comparable number of significant differentially phosphorylated sites were found in both starvation conditions; in wildtype cells 46 phosphorylation sites were increased in abundance when starved for carbon and 49 were increased in abundance when starved for phosphorus (**Figure 4D)**. GSEA analysis revealed that protein phosphorylation events are enriched for multiple processes related to translation in in both starvation conditions (**Figure 4E**) consistent with the functional targets of RIM15 being primarily related to translational activity and protein homeostasis in quiescence (Sun et al. 2020).

Phosphorylation events on 11 different proteins were found to be commonly regulated by RIM15 in both starvation conditions (p.adj < 0.01, hypergeometric test) (**Figure 4D**). The change (expressed as a log_2_ fold change) of each phosphorylation event between WT and *RIM15Δ0* cells shows a clear gradual up-regulation with time (**Figure 4F**), which we confirmed is not a result of increased protein levels (**Figure S4B**). Common phosphorylation events are found in proteins involved in three key biological processes: 1) actin polymerization (n=2, ABP1 (Ser165) and BZZ1 (Ser472), 2) glycogen metabolism (n=2 YPI1 (Ser133) and GLC8 (Ser184)), and 3) proteostasis (n=7). RIM15 protein phosphorylation targets involved in translation include the translation initiation factor TIF4631 (Ser163), ribosome proteins RPL7A (Ser8) and RPL7B (Ser8), STM1 (Ser118) which is required for optimal translation under nutrient stress, RPC19 (Ser15) the common subunit of RNA polymerases I and III, DPS1 (Ser14) a tRNA synthetase, and YRB1 (Ser60) involved in ubiquitin-mediated protein degradation during the G1/S transition of the cell cycle. We searched for known protein-protein relationships between these common targets of RIM15 using the STRING database (Szklarczyk et al. 2021). Although some proteins are known to have functional relationships with each other, none of those proteins has previously been identified as being a direct target of RIM15 (**Figure 4G**).

To test the essentiality of protein phosphorylation targets of RIM15 in quiescence, we examined the survival of the corresponding gene deletion mutant in starvation conditions (Sun et al. 2020). Overall, 20 (carbon starvation) and 30 (phosphorus starvation) genes and proteins were common to both studies. Not all RIM15 protein phosphorylation targets have reduced survival in quiescence when deletion (**Figure S4C**). A total of 6/20 (carbon starvation) and 17/30 (phosphorus starvation) have a reduced starvation survival when deleted, with the extreme cases being SNF1 and IGO1, which are both phosphorylation targets of RIM15 that have strongly reduced survival in the absence of RIM15 in carbon and phosphorus starvation respectively.

### Phosphoproteome profiling identifies condition dependent signaling pathways

To study the dynamics of the phosphoproteome, we applied WGCNA (**Figure S5A)**. In total, we identified 1,472 unique phosphorylated proteins, out of which 1,340 phosphorylation events were differentially expressed as a function of genotype, nutrient starvation, or time (**Table S2**). WGCNA co-expression clustering of phosphorylation events identified eight Phosphorylation Expression Modules (PEMs) (**Figure S5B**), and functional enrichment was performed for each cluster (**Figure S5C**). Hundreds of protein phosphorylation events are reduced upon carbon removal (e.g. PEM1, PEM4, PEM6, PEM7) and phosphorus starvation (PEM1, PEM5, PEM7) (**Figure S5D)**. The dynamics of the majority of phosphorylation events differs between starvation conditions as PEM1 (n = 452) is the only module for which the dynamics are largely consistent between conditions. The corresponding proteins in PEM1 are enriched for functions in the cell cycle and MAPK signaling pathways, translation initiation factor binding, histone binding, and enzyme activities (**Figure S5C**). However, overall these results indicate pervasive protein phosphorylation independent of RIM15 during the initiation of quiescence in response to carbon and phosphorus starvation (**Figure S5E)**.

As with proteome remodeling, most protein phosphorylation events during quiescence establishment vary depending on the starvation signals. For example, phosphorylated proteins in PEM2 and PEM3 share similar dynamics in carbon starvation conditions but have distinct patterns in phosphorus starvation. The differing dynamics of PEM2 and PEM3 suggests that phosphorylation of proteins involved in cytoskeleton organization is increased in carbon starvation but not drastically remodeled under phosphorus starvation. PEM4, which is enriched for translation initiation functions, is continuously downregulated in carbon starvation. Similarly, PEM5 and PEM7, which are enriched for autophagy and enzyme functions (**Figure S5C)**, are systematically downregulated in phosphorus starvation (**Figure S5D**). Some phosphorylation dynamics are found in response to both starvation conditions and are consistent with known processes in quiescence. For example, PEM8 contains protein phosphorylation targets that rapidly increase following carbon or phosphorus starvation. PEM8 is enriched for proteins regulating nuclear chromatin including SWI/SNF complex, which is in line with the proposed mechanism of chromatin condensation in repressing transcription globally during quiescence (Li, Miles, and Breeden 2015; Spain, Braceros, and Tsukiyama 2018).

In contrast to proteome dynamics, the phosphoproteome appears to continue changing well beyond the initiation of quiescence as significant changes in protein phosphorylation levels are observed between 16 hours and 30 hours. This suggests that signaling pathways actively regulate processes in quiescent cells long after the starvation signal is first sensed.

## Discussion

In this study, we undertook a parallel analysis of the yeast proteome and phosphoproteome during the initiation of quiescence in response to two distinct nutrient starvations: carbon and phosphorus. To define the role of RIM15 we studied both wildtype cells and an isogenic *RIM15Δ0* strain. Our analysis of expression of 1,277 proteins and 1,472 phosphorylation events corresponding to 785 phosphoproteins reveals that proteome and phosphoproteome remodeling in quiescent cells is largely dependent on the specific starvation signal. We also find distinct behaviors with respect to the dynamics of expression and phosphorylation. Whereas the major changes in the proteome occur within 6 hours of starvation, the phosphoproteome is continuously remodeled for as long as 30 hours following the starvation signal.

Our study confirms previous observations regarding the increased expression of mitochondrial proteins in quiescence under carbon starvation (Davidson et al. 2011). Interestingly, we find that these proteins are also up-regulated in phosphorus starvation suggesting that increased mitochondrial activity is essential for initiation of quiescence independent of starvation signal. Our results are also consistent with a previous study of amino acid starvation that found that a successful starvation response is correlated with expression of genes encoding oxidative stress response and mitochondrial functions regardless of the starvation nutrient (Petti et al. 2011). Mitochondria undergo substantial reorganization during quiescence entry (Laporte et al. 2018). Understanding the causal relationships between mitochondrial remodeling and increased mitochondrial protein expression is an important area for further investigation.

Previously, genetic interaction analysis suggested new roles for the serine/threonine kinase RIM15 in quiescence involving both transcription and translation (Sun et al. 2020). By looking at the overall trend of differential phosphorylation events between wildtype and *RIM15Δ0* cells, we found additional evidence for RIM15 in regulating protein homeostasis via phosphorylation of proteins involved in translation and amino acid metabolism. This includes processes involved in RNA processing (including rRNA, ncRNA and mRNA) and translation. Surprisingly, we do not find evidence for widespread protein phosphorylation by RIM15 that is specific to quiescence. This is likely due to two reasons. First, we find that protein phosphorylation by RIM15 occurs in actively proliferating cells as exemplified by phosphorylation of Serine 64 on IGO1. Second, there is likely significant functional redundancy among protein kinases that function during quiescence.

Our study demonstrates the feasibility of using SILAC with prototrophic yeast strains in both rich and minimal starvation conditions. However, we identified a key limitation in the use of SILAC for labeling nitrogen starved cells. Our mass spectrometry analysis demonstrates that in nitrogen starvation media supplemented labeled amino acids are degraded and incorporated into newly synthesized amino acids resulting in unanticipated spectra. Thus, in addition to the activation of autophagy in response to nitrogen starvation (Tyler and Johnson 2018), cells appear to catabolize amino acids and activate salvage pathways for amino acid synthesis. The role of amino acid catabolism in quiescent cells starved for nitrogen warrants further investigation.

## Methods

### Strains, Culture Conditions, and SILAC Labeling

All experiments were performed using S288c isogenic strains. For label incorporation tests we used the prototrophic haploid strain FY4 (*MATa*), an auxotrophic strain derived from FY4 (*MATa* lys2Δ0 arg6Δ0) and a prototrophic *RIM15Δ0* (*MATa RIM15Δ0*::kanMX) created from FY4. Deletion of *RIM15* was confirmed by PCR. For all subsequent proteomic and phosphoproteomic experiments prototrophic FY4 (*MATa*) and *RIM15Δ0* (*MATa RIM15Δ0*::kanMX) were used. SILAC experiments for incorporation tests were performed in cells grown in synthetic complete (SC) medium lacking arginine and lysine. SILAC media was prepared by supplementing with light, medium or heavy isotopes of arginine and lysine (L-lysine/L-arginine, 4,4,5,5-D4 L-lysine/^13^C_6_-arginine, ^13^C_6_^15^N_2_-lysine/^13^C_6_^15^N_4_-arginine) (Sigma). For quantitative SILAC experiments in starvation conditions, cells were first cultured in SILAC medium supplemented with either of the three isotope labeled medium for 10 generations for maximal isotope incorporation and then cells were pelleted and washed twice with water before transferring into the corresponding isotope supplemented medium. Cells labeled in SILAC medium were split into three equal portions and inoculated into 300mL of nutrient depleted medium lacking either ammonia (N, 0mM nitrogen), phosphorus (P, 0mM phosphorus), or very low carbon (C, 3.35mM glucose), with an initial concentration of 2 x 10^6^ cells/mL.

For viability quantification at each time point, 1 × 10^7^ cells were collected and subsequently washed once with sterilized DI water and one more time with PBS. The washed cell pellet was resuspended with 1mL of 1x PBS and stained with 3.34µM of SYTO® 9 and 20µM of propidium iodide for 20 minutes. The stained samples were then analyzed by flow cytometry (BD Accuri^TM^ C6).

To quantify the proteome and phosphoproteome at multiple timepoints during quiescence in response to three nutritional starvations, we performed three independent experiments for each genotype (WT and *RIM15Δ0*) in each condition. We collected three time points after transferring the sample to nutrient depleted medium at T=6h, T=16h, and T=30h. T0 samples were collected from the SILAC medium right before cell transfer.

### Protein extraction, pooling and digestion

At each time point 3 x 10^8^ cells were pelleted and washed twice with ice cold PBS before flash freezing. A technical replica was also collected. Samples were randomized for protein extraction and digestion to minimize technical variation. After thawing on ice, cells were disrupted by glass-bead agitation at 4°C in standard buffer: 50 mM Tris-HCl pH 7.5, 150 mM NaCl, 20 mM iodoacetamide (IAM), 1x complete protease inhibitor (Roche) and 1x phosphatase inhibitor cocktail (PhosphoSTOP, Roche). The extract was cleared by centrifugation and protein concentration was determined by Bradford assay (BioRad). Cell lysate extracted from different genotypes under the same starvation treatment at the same time point were then mixed at equal amounts with an external control sample (wild type cells in exponential growth phase in rich medium) at a 1:1:1 ratio. Approximately 600μg of the mixed light/medium/heavy/ protein sample was processed for in-solution digestion as described in (Monteoliva et al. 2011). Proteins were reduced with 5mM DL-Dithiothreitol (DTT) for 30 minutes at 37°C and alkylated with 10 mM iodoacetamide for 30 min at 30°C. Samples were diluted six times with 25 mM ammonium bicarbonate, trypsin/lysC (Promega, Spain) was added to the protein mixture at a 25:1 protein:protease ratio (w/w), and samples were incubated overnight at 37°C. Protein digestion was stopped by addition of formic acid.

### Phosphopeptide enrichment

Enrichment for phosphopeptide was performed using Pierce™ TiO2 Phosphopeptide Enrichment and Clean-up Kit. All digested peptides were desalted with C18 according to a previous study (Villén and Gygi 2008). Then, an aliquot of 10 μg peptides was separated to be further processed and analyzed without phosphopeptide enrichment. The remainder of the sample was processed with the TiO2 according to the user manual. Samples representing the whole proteome were solubilized in 15 μl of 2% ACN 0.5% AcOH and 2 μl were analyzed by LC-MS/MS. Enriched phosphopeptides were solubilized in 10 μl of 2% acetonitrile 0.5% acetic acid and 5 μl were analyzed by LC-MS/MS.

### Mass Spectrometry

LC separation was performed online on EASY-nLC 1000 (Thermo Scientific) using Acclaim PepMap 100 (75 um x 2 cm) precolumn and PepMap RSLC C18 (2 μm, 100A x 50 cm) analytical column. Peptides were gradient eluted from the column directly to Orbitrap HFX mass spectrometer using 160 min acetonitrile gradient from 5-26% B in 118 minutes followed by ramp to 40% B in 20 min and final equilibration in 100% B for 15 minutes (A=2% acetonitrile 0.5% acetic acid; B=80% acetonitrile 0.5% acetic acid). The flow rate was set at 200 nl/minute.

For whole cell digests, high resolution full MS spectra were acquired with a resolution of 120,000, an AGC target of 3e6, with a maximum ion injection time of 32 ms, and scan range of 400 to 1600 m/z. Following each full MS scan 20 data-dependent HCD MS/MS scans were acquired at the resolution of 7,500, AGC target of 2e5, maximum ion time of 32 ms, one microscan, 1.4 m/z isolation window, nce of 27 and dynamic exclusion for 45 seconds.

For phosphopeptides analysis, high resolution full MS spectra were acquired with a resolution of 120,000, an AGC target of 3e6, with a maximum ion injection time of 100 ms, and scan range of 400 to 1600 m/z. Following each full MS scan 20 data-dependent HCD MS/MS scans were acquired at the resolution of 30,000, AGC target of 5e5, maximum ion time of 100 ms, one microscan, 1.4 m/z isolation window, nce of 27 and dynamic exclusion for 45 seconds.

### Protein identification

MS data were analyzed using MaxQuant software version 1.6.3.4 and searched against the *S. Cerevisiae* reference database (http://www.uniprot.org/) containing 6,721 entries. Multiplicity was set to three, matching the number of SILAC labels used (“light,” “medium,” and “heavy”) in each experiment; lys-4/arg-6 and lys-8/arg-10 were specified as medium and heavy labels, respectively. Database search was performed in Andromeda integrated in the MaxQuant environment. A list of 248 common laboratory contaminants included in MaxQuant was also added to the database as well as reversed versions of all sequences. For searching, the enzyme specificity was set to trypsin with the maximum number of missed cleavages set to 2. The precursor mass tolerance was set to 20 ppm for the first search used for non-linear mass re-calibration and then set to 6 ppm for the main search. Oxidation of methionine was searched as variable modification; carbamidomethylation of cysteines was searched as a fixed modification. For phosphopeptides samples phosphorylation of serine, threonine and tyrosine residues was also included as a variable modification. The false discovery rate (FDR) for peptide, protein, and site identification was set to 1%, the minimum peptide length was set to 6. To transfer identifications across different runs, the ‘match between runs’ option in MaxQuant was enabled with a retention time tolerance of one minute (after data-dependent nonlinear retention time recalibration).

### Data analysis

Data analysis was performed using R. The triplicate SILAC experiments were combined to get a ratio for each protein and phosphorylation event across the four time points relative to the “light labeled” internal control. The median normalized ratios were used for downstream analysis. Only proteins and phosphorylation events that had valid values for which over 75% of the samples passed filter (including 3 biological replicates, 2 genotypes, 2 nutritional conditions and 4 timepoints) were considered for further quantitative analysis.

#### Differential expression analysis of proteome and phosphoproteome

Differential expression was defined by identifying proteins/peptides with between time point, genotype, nutrient variance significantly larger than within-replicate variance using ANCOVA (analysis of covariance). Specifically, we applied a two-step procedure to define DE events as follows: (i) calculate F statistic and P values (defined as the ratio of between-group variance relative to within-group variance) for each protein. The P value distributions were skewed towards 0, indicating a large number of DE proteins and peptides during quiescence entry (ii) correct P values for multiple testing using the Benjamini-Hochberg (BH) method. We selected DE proteins and phosphorylation events by applying a threshold of corrected p-value of0.05.

#### Weighted gene co-expression network analysis (WGCNA) clustering analysis

WGCNA analysis (Zhang and Horvath 2005) was carried out using the WGCNA R package (Langfelder and Horvath 2008). To define whole proteome co-expression modules (WPMs), only the 1,067 DE proteins were considered. A Pearson correlation matrix (with direction, i.e. for building signed co-expression network) was calculated using DE proteins across the 42 samples and an adjacency matrix was calculated by raising the correlation matrix to a power of 6 using the scale-free topology criterion (Zhang and Horvath 2005). Co-expression modules were defined by hybrid dynamic tree-cutting method (Langfelder, Zhang, and Horvath 2008) with the minimum height for merging modules at 0.25. For each co-expression module, a consensus trend was calculated based on the first principal component (also known as eigengene) and cluster membership was defined by the Pearson correlation between an individual protein and the consensus of the WPM. Proteins were assigned to the most correlated co-expression cluster (i.e. WPM) with a cutoff of r ≤ 0.7. Individual WPMs were annotated using two pathway databases (Gene Ontology and KEGG) using Fisher’s exact test and applying a FDR ≤ 0.05). Phosphoproteome co-expression modules (PEMs) were also defined by the same procedure but with an adjacency matrix calculated by raising the correlation matrix to a power of 8 using the scale-free topology criterion.

#### Differential expression analysis using ANCOVA

To model each protein or phosphorylation event within a starvation condition, we used the normalized ratio at different time points as the dependent variable with the two different genotypes (WT and *RIM15Δ0*) treated as independent categorical variables, and sampling time as the covariate. We modeled each protein or phosphorylation event using ANCOVA as follows:

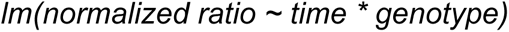

Where *time* is the hours post nutrient removal (0hr, 6hr, 16hr, 30hr), and *genotype* is either WT or *RIM15Δ0.* We corrected p-values for multiple testing by applying the Benjamini-Hochberg (BH) method, and then selected DE proteins and phosphorylation events by applying a threshold of corrected p-value ≤ 0.1.

Coefficients from ANCOVA were then subject to Gene Set Enrichment Analysis using *clusterProfiler()* package in R (Yu et al. 2012).

## Acknowledgements

We thank the members of the Gresham lab and Vogel lab for helpful discussions. The mass spectrometric experiments were supported in part by NYU Langone Health. This work was supported by the NIH (R01 GM107466).

## Authorship contributions

SS and DG designed the study; SS performed the experiments; SS analyzed the data in consultation with DT; SS wrote the first draft and DG and SS revised the paper.

## Data availability

Data are available via ProteomeXchange with identifier PXD044239.

## Supplemental figures

**Figure S1.**
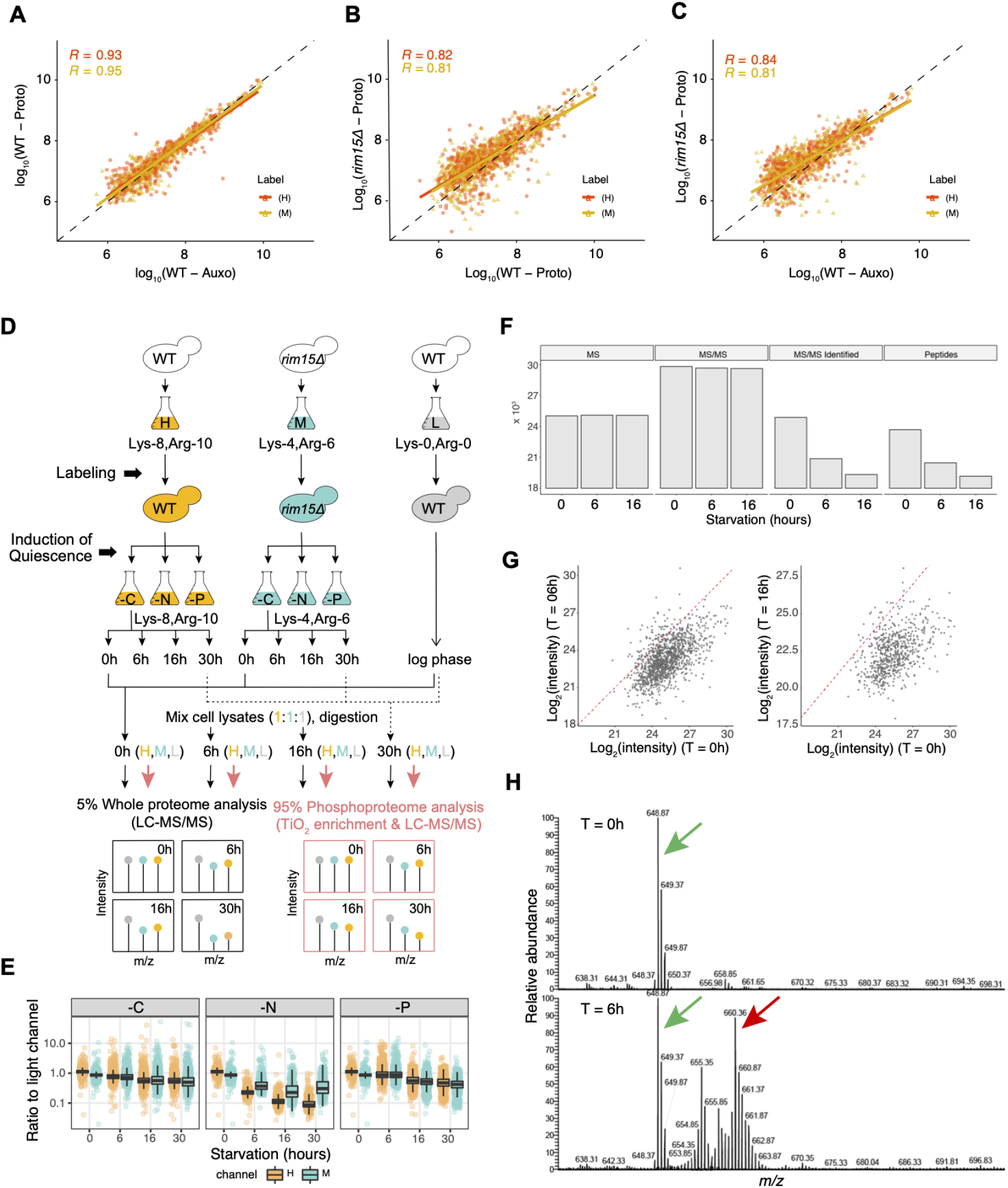
Whole proteome comparison between SILAC studies using different genotypes. Comparison of protein abundances identified in **A)** wildtype auxotrophic and prototrophic cells **B)** wildtype prototrophic and *RIM15Δ0* prototrophic cells and **C)** wildtype auxotrophic and *RIM15Δ0* prototrophic cells grown in the presence of either heavy (H) or medium (M) isotope medium. Pearson correlations are indicated using the same color.**D)** Experimental design for simultaneous profiling of proteome and phosphoproteome during quiescence initiation. **E)** Ratios of intensities from heavy and medium channels (staved cells) to the light channel (log phase cells) show marked signal reduction in cells initiating quiescence in response to nitrogen starvation. **F)** Comparison of different LC/MS characteristics across timepoints in cells starved for nitrogen. **G)** Comparison of log_2_(intensity) of peptides detected using mass spectrometry at different timepoints following nitrogen starvation. **H)** MS/MS analysis on heavy isotope labeled cells collected T=0h and T=6h. A representative spectra of a major peptide peak (green arrow) detected at T=0h. Whereas the peak is also detected in a sample 6 hours after nitrogen starvation, a novel second peak (red arrow) is apparent in the sample. This additional unexpected peak is consistent with cells metabolizing lys-8/arg-10 and incorporating heavy isotopes into other amino acids.

**Figure S2.**
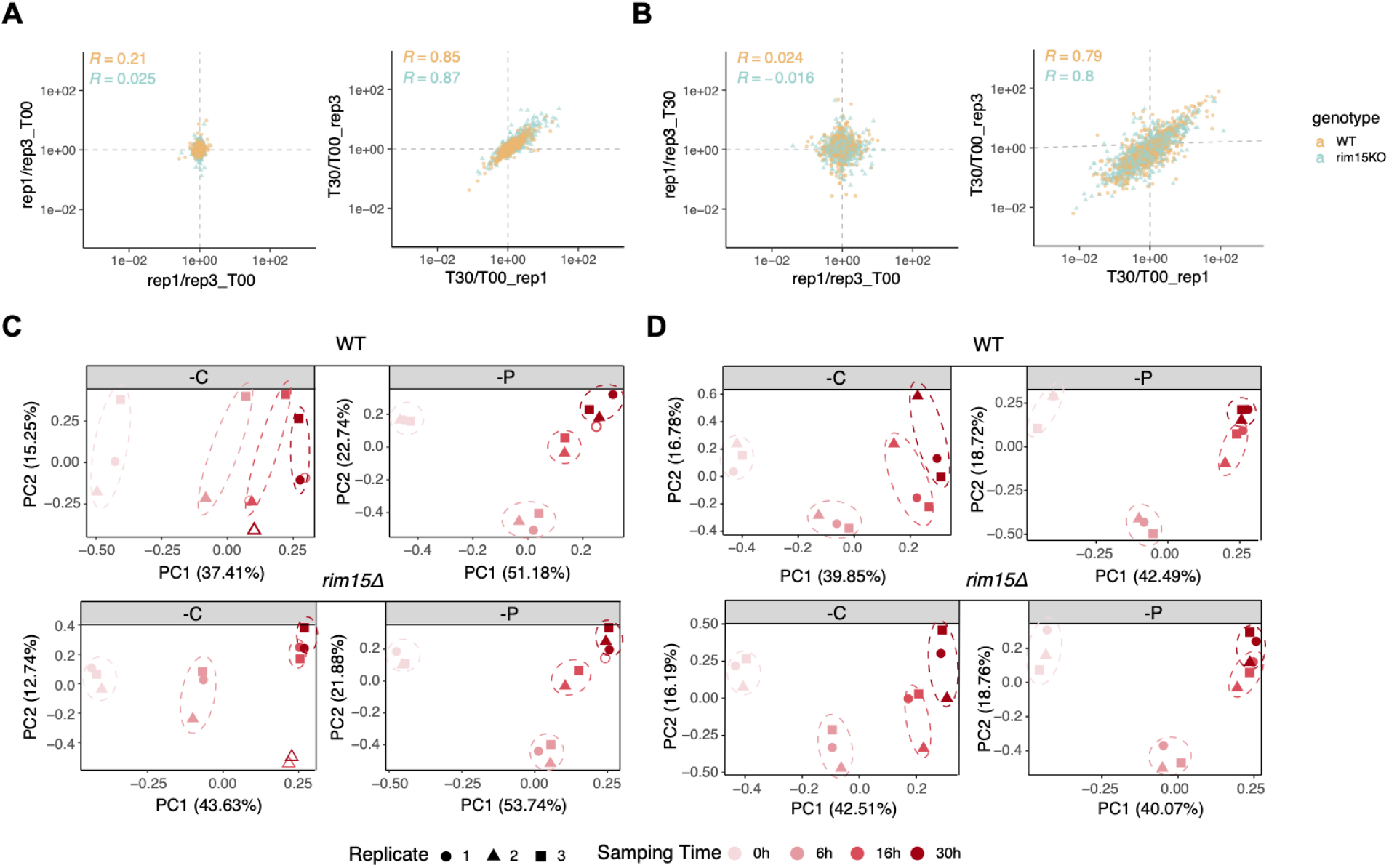
Sources of variation in proteome and phosphoproteome data. Representative null comparisons (0h_replicate 1/0h_replicate 3 and 30h_replicate 1/30h_replicate 3) display markedly distinct patterns from true comparisons (30h_replicate 1/0h_replicate 1 and 30h_replicate 3/0h_replicate 3) for whole proteome (**A**) and whole phosphoproteome (**B**) data for both genotypes. Correlation coefficients (r) for each comparison is indicated. Principal-component analysis of whole proteome data (**C**) and phosphoproteome data (**D**). Biological replicates from different time points (red gradient color) post nutrient starvation cluster together and are dispersed largely along the first principal component in both wildtype (WT) and *RIM15Δ0* cells.

**Figure S3.**
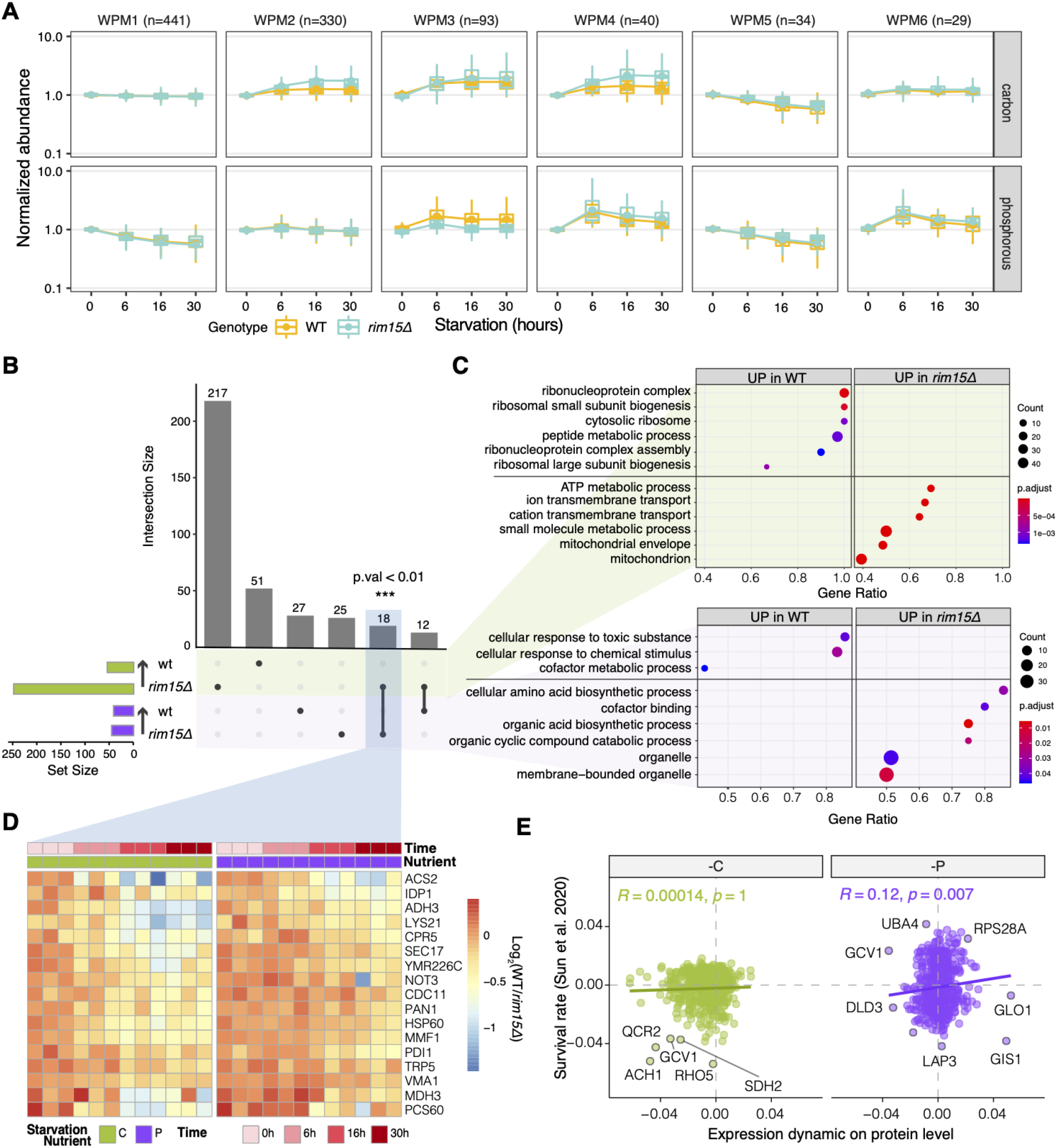
Functional impact of RIM15 on global protein expression. **A)** Proteome dynamics of whole proteome modules (WPMs) for WT and *RIM15Δ0* cells separated by starvation conditions. **B)** Summary of significantly differentially expressed proteins in WT and *RIM15Δ0* as a function of time of starvation identified using ANCOVA. A single protein (excluded for plotting purposes) was found to be upregulated in wild type cells in both conditions, SBP1, which binds to eIF2 and represses translation and forms P bodies by binding to mRNAs under carbon starvation (Segal, Dunckley, and Parker 2007). **C)** GSEA of significant DE proteins between WT and *RIM15Δ0* in response to carbon (green) and phosphorus (purple) starvation. **D)** Differential expression log_2_(WT/*RIM15Δ0*) of 18 commonly up regulated proteins in *RIM15Δ0* in both starvation conditions. **E)** Comparison between survival rate (y-axis) determined using gene deletion analysis (Sun et al. 2020) and the dynamics of differentially expressed proteins (x-axis). Pearson correlation is indicated on the plot.

**Figure S4.**
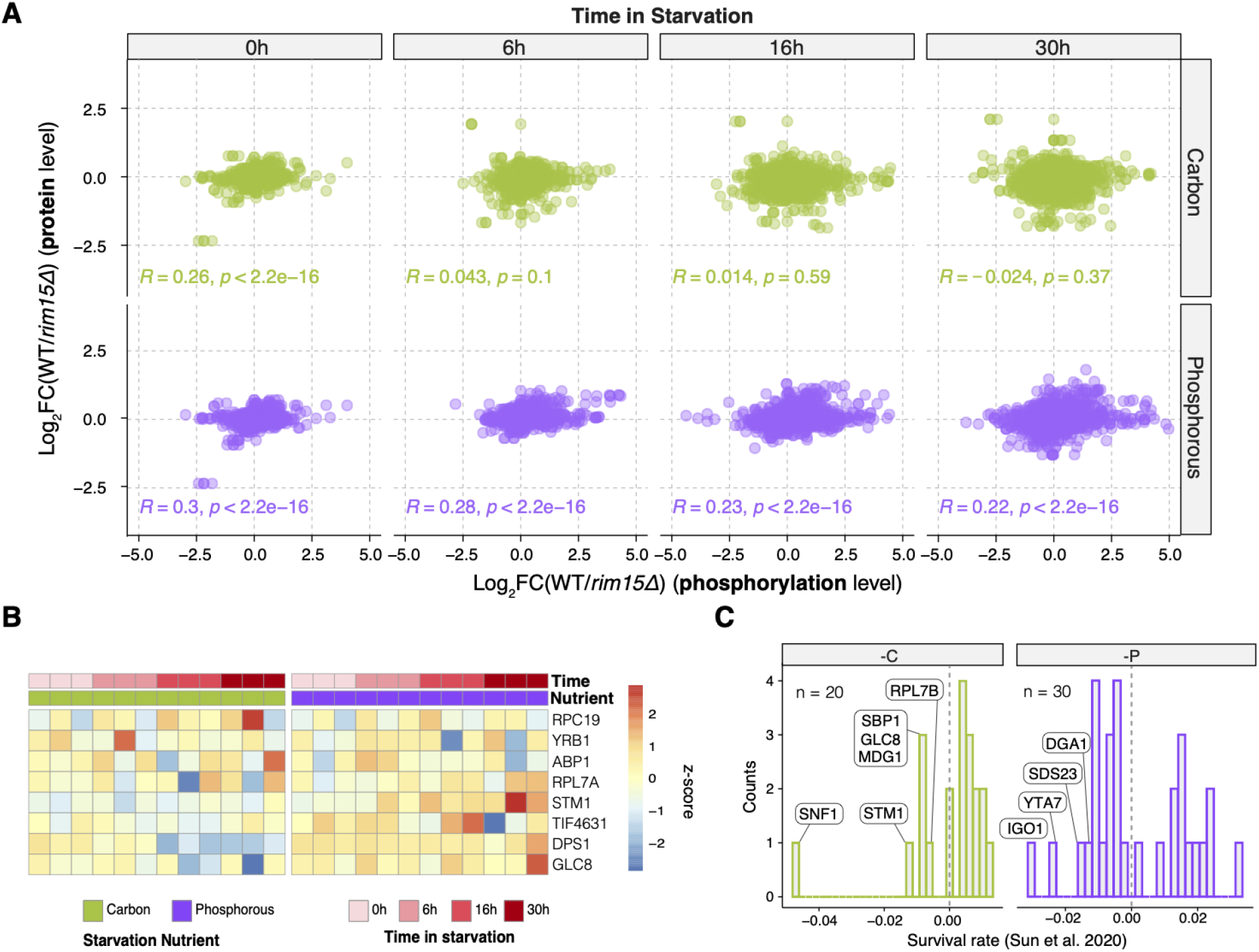
Functional impact of RIM15 on protein phosphorylation events during entry of quiescence. **A)** Fold changes of protein phosphorylation state compared with fold change in protein expression level for WT versus *RIM15* null mutants. **B)** Protein expression levels of proteins whose phosphorylation is regulated by RIM15 in both starvation conditions. For visualization purposes data are scaled by rows. **C)** Viability of gene deletion mutants in carbon or phosphorous starvation conditions for putative RIM15 phosphorylation targets (Sun et al. 2020).

**Figure S5.**
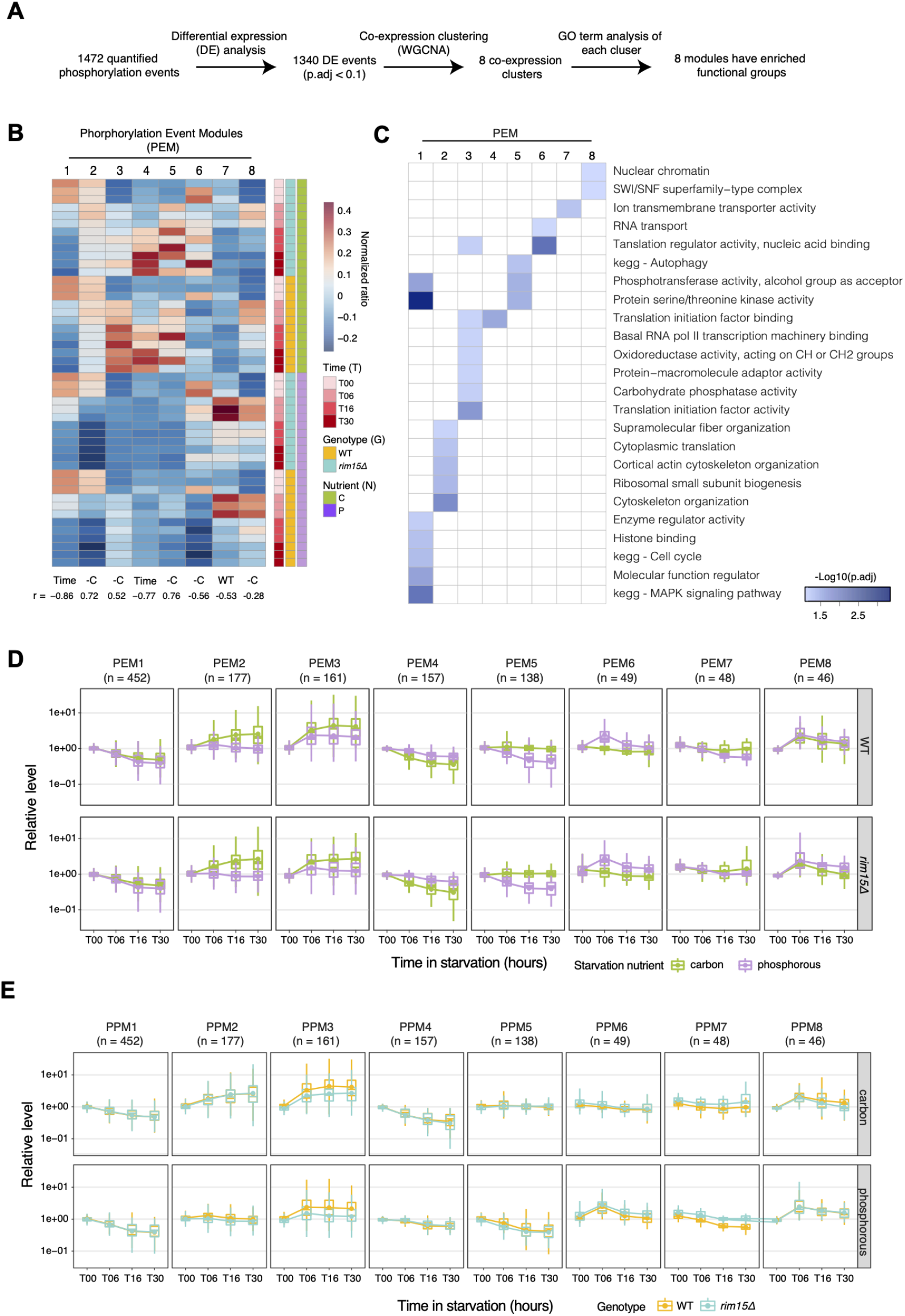
Phosphoproteome profiling reveals coordinated protein phosphorylation events. **A)** Overview of computational analysis of the phosphoproteome. **B)** Mean expression of consensus eignegenes for each phosphorylation expression module (PEM). Each column indicates a module and each row is its expression in a single sample. **C)** Functional annotations of PEMs using GO and KEGG annotations (FDR < 0.05). **D)** Dynamics of phosphorylation events in eight PEMs separated by genotype and colored by starvation nutrients. **E)** Dynamics of phosphorylation events in eight PEMs separated by starvation condition and colored by genotype.

## Supplemental Tables

**Table S1.** Abundance of 1,277 proteins detected in WT and *RIM15Δ0* cells starved for glucose of phosphorus.

**Table S2.** Abundance of 1,472 unique phosphorylated proteins detected in WT and *RIM15Δ0* cells starved for glucose of phosphorus.

**Table S3.** Identity of 11 proteins that are upregulated in carbon starvation (p.adj < 0.05) are genetically required for survival (p.adj < 0.05)

**Table S4.** Identity of 13 proteins that are upregulated in carbon starvation (p.adj < 0.05) are genetically required for survival (p.adj < 0.05)

